# Pan-cortical cellular imaging in freely behaving mice using a miniaturized micro-camera array microscope (mini-MCAM)

**DOI:** 10.1101/2024.07.04.601964

**Authors:** Jia Hu, Arun Cherkkil, Daniel A. Surinach, Ibrahim Oladepo, Ridwan Hossain, Skylar Fausner, Kapil Saxena, Eunsong Ko, Ryan Peters, Michael Feldkamp, Pavan C Konda, Vinayak Pathak, Roarke Horstmeyer, Suhasa B Kodandaramaiah

**Author notes:** Corresponding author Send Correspondence to: Suhasa B Kodandaramaiah Associate Professor, Department of Mechanical Engineering, University of Minnesota, Twin Cities, MN 111 Church St SE, Minneapolis, MN – 55455. Equal contribution (Jia Hu, Arun Cherkkil).

## Abstract

Understanding how circuits in the brain simultaneously coordinate their activity to mediate complex ethnologically relevant behaviors requires recording neural activities from distributed populations of neurons in freely behaving animals. Current miniaturized imaging microscopes are typically limited to imaging a relatively small field of view, precluding the measurement of neural activities across multiple brain regions. Here we present a miniaturized micro-camera array microscope (mini-MCAM) that consists of four fluorescence imaging micro-cameras, each capable of capturing neural activity across a 4.5 mm x 2.55 mm field of view (FOV). Cumulatively, the mini-MCAM images over 30 mm^2^ area of sparsely expressed GCaMP6s neurons distributed throughout the dorsal cortex, in regions including the primary and secondary motor, somatosensory, visual, retrosplenial, and association cortices across both hemispheres. We demonstrate cortex-wide cellular resolution *in vivo Calcium (Ca^2+^)* imaging using the mini-MCAM in both head-fixed and freely behaving mice.

## INTRODUCTION

Neuronal activity distributed across multiple regions of the brain mediates behavior. Recent advances in simultaneous large-scale neural recording across multiple cortical regions have revealed crucial insights into how the neocortex is recruited in a wide variety of sensory-motor^1^ and cognitive tasks^2^. Behavioral states, attention, motor activity, and state of arousal have dramatic effects on spontaneous activity across the cortex^3–5^. These distributed activities are altered during learning^6,7^ and in diseased states^8^, and influence how the underlying neurons within these circuits respond to evoked sensory stimuli^9^. Thus, it is crucial to simultaneously record from multiple cortical regions during behavior.

Much of the work studying large-scale distributed activity across the cortex has been done under head fixation. Instrumentation capable of imaging single neuron activities across several regions of the cortex using single photon(1P)^10–12^ and two photon (2P) imaging modalities^13–16^ have been developed. This has been achieved by scaling up the size of the objective and corresponding optical elements associated with them. The size and weight of such conventional optical elements need to increase cubically with increased FOV, if the same imaging resolution is desired. This makes such scaling incompatible with miniaturization for imaging in freely behaving animals. Thus far even highly optimized miniaturized imaging devices^17^ have been limited to imaging < 4mm diameter area in mice. While this allows imaging across a few functional distinct regions, it precludes investigations of how activity is globally coordinated across the cortex in complex behaviors such as spatial navigation^18^. While alternate approaches such as a lensless fluorescence imaging^19–21^ and computational miniaturized microscopes have been developed^22^ , practical demonstration of cellular resolution imaging with such devices in freely behaving mice has not yet been realized.

We bring together two innovations to realize a miniaturized technology that can reliably image 32 mm^2^ of the dorsal cortical surface. First, we developed a multiplanar faceted cranial window that allows optical access to 52 mm^2^ of the cortex, while creating four individual planar tissue surfaces which can be imaged using 1 photon (1P) imaging optics. Second, we developed a compatible miniaturized micro-camera array microscope (mini-MCAM), that consists of four micro-cameras, each capable of fluorescence imaging of 4.5 mm x 2.55 mm FOVs. Cumulatively, the mini-MCAM images several cortical regions simultaneously in sparsely expressed GCaMP6s cells distributed throughout the cortex, including the primary and secondary motor, the somatosensory, the visual, retrosplenial and association cortices across both hemispheres. We demonstrate the capability of the mini-MCAM to capture robust Calcium (Ca^2+^) signals from individual neurons in both headfixed and freely behaving mice. The mini-MCAM can be easily assembled using off-the-shelf miniaturized optical imaging modules and provides a novel solution for cortex-wide imaging of cellular activities in freely behaving mice.

## RESULTS

### Multi planar-faceted cranial window for accessing > 50 mm^2^ for 1P optical imaging

We aimed to image large areas of the dorsal cortex up to a depth of 200 μm using the mini-MCAM. This poses two challenges. First, optics needed to image large FOVs while maintaining sufficient resolution to distinguish individual cellular elements. If traditional multi-element lenses are scaled to increase FOV, this results in cubic scaling of both size and weight of the optics, while incompatible with miniaturization (**Fig 1A left**). An alternative approach used in this work is to use arrays of micro-cameras that are designed for small FOVs and cumulative images of a large area of the brain surface (**Fig. 1A right**). A second challenge to contend with is the curvature of the brain. Traditionally, 1P cortical imaging has been done via flat cranial windows^23^. For imaging the whole dorsal cortex, or the whole cerebellar cortex, curved cranial windows have been developed^24–27^. However, imaging curved surfaces limits the overall area of the optically accessible surface (**Fig. 1B**). To solve this limitation, we engineered a cranial window that comprises multiple planar facets that make large areas of the bilateral cortex optically accessible by creating flat tissue planes that can be imaged using the mini-MCAM, while also minimizing brain displacement underneath the implant. This implant consists of a 3D printed frame, to which a PET film creased to form four planar facets is bonded. The implant additionally has a titanium headplate to allow head fixation of the animal if required (**Fig. 1C**). The multiplanar faceted cranial windows could be chronically implanted for long durations, providing optically clear access to 52 mm^2^ of the dorsal cortex (**Fig. 1D**).

**Figure 1:**
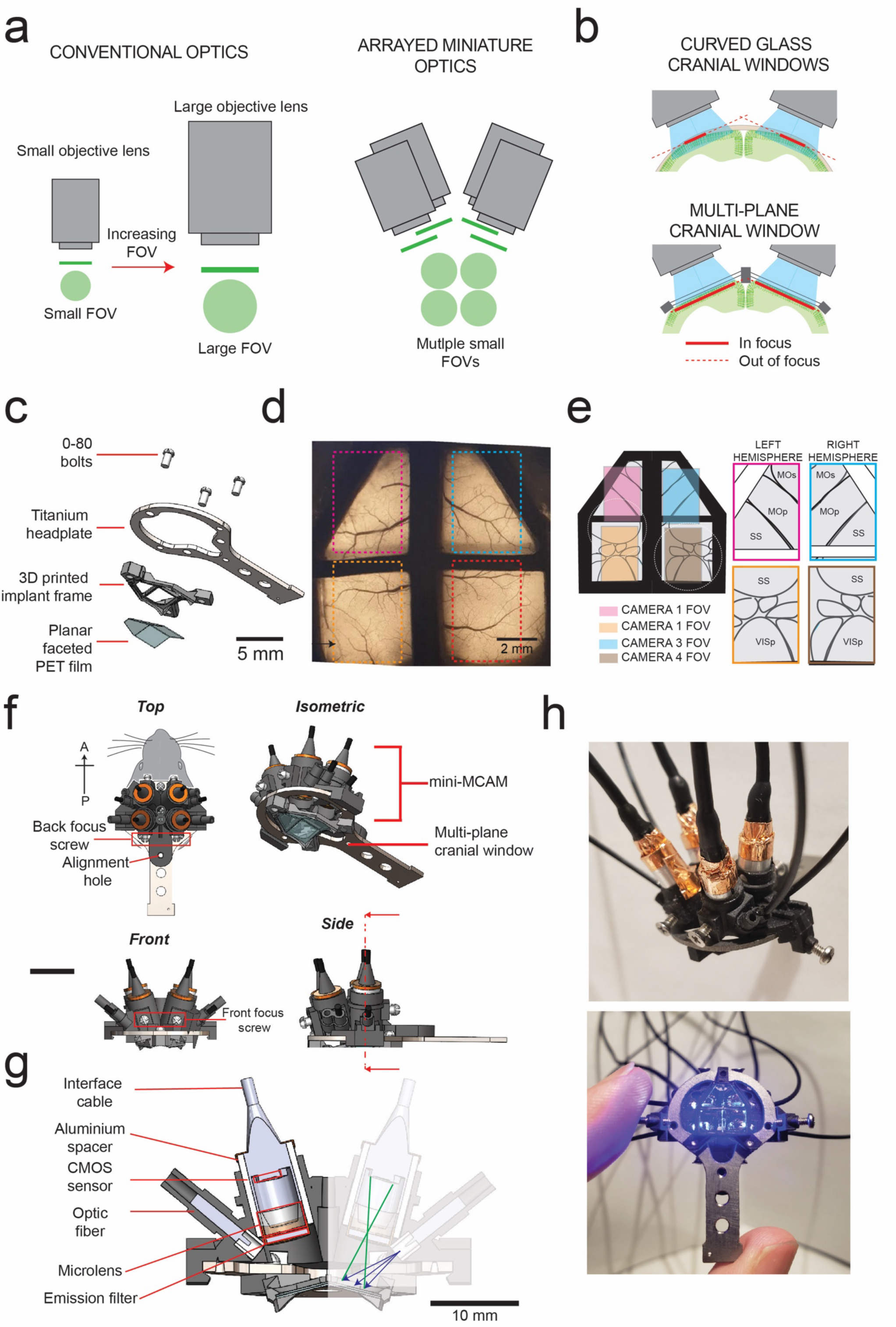
mini-MCAM for simultaneous multi-site cellular resolution imaging across the dozens of cortical regions: (a) Concept of arrayed miniaturized optics for imaging large areas – *left:* scaling up conventional multi-element optics for imaging larger field of views (FOVs) at the same resolution results in cubic scaling of optical elements. Right: Using arrays of miniaturized imaging elements results in linear scale of size and weight. (b) Imaging the brain through curved cranial windows (top) implanted across the dorsal cortex results in significant reduction of tissue area in focus. Bottom: Multiplane cranial window allows both optical access to bilateral cortex, while creating planar tissue facets for imaging with 1-photon imaging optics. (c) Computer aided design (CAD) schematic of the multi-planar faceted cranial window. (d) Widefield epi-fluorescence imaging of a Cux2-CRE-ERT2. x Ai162 double transgenic mouse chronically implanted with the multi-planar faceted image shown in **c**. Dashed rectangles indicate regions of cortex imaged using each of the micro-cameras in the miniaturized micro-camera array microscope (mini-MCAM) shown in **f-h**. (e) Schematic of the different cortical areas imaged simultaneously using the mini-MCAM. (f) CAD schematic of the isometric, top, front and side views of the mini-MCAM. Scale bars indicate 5 mm. (g) Cross-sectional view of the internal elements mini-MCAM. Cross section taken at the plan indicated in the side view schematic in (f). (h) Photographs of the mini-MCAM. Scale bars indicate 5 mm

### Mini-MCAM design

The mini-MCAM consists of an array of 4 individual micro-cameras. Each micro-camera contains its own CMOS sensor and miniaturized imaging optics (lens assembly, optical filters, and illumination sources) that stream microscopic video data simultaneously. The micro-cameras are installed within a main housing that is mounted on the frame of an implanted cranial window (**Fig. 1F**).

Two micro-cameras are positioned to image bilateral regions of the somatosensory and visual cortices and two micro-cameras to image the frontal cortex bilaterally. A custom-diced 3 mm diameter emission filter is placed outside the lens. Focusing is achieved manually by adjusting micro-camera placement inside the frame of the 3D printed scope housing. Four 250 μm optical fibers coupled to a diode laser provide sufficient wide-field excitation illumination to each FOV (461.3 ± 3.3 nm, ∼3.3-4.2 mW)^28^ . The mechanical housing holds each micro-camera aligned to the normal of each planar face of the cranial window surface. The sensors are adapted from micro-cameras utilized for endoscopic imaging and have all the electronic readout circuitry placed behind each micro-camera CMOS sensor, rather than adjacent to, as in most CMOS sensor layouts, which is critical for minimizing each unit’s total lateral footprint. A small footprint increases the micro-camera packing density and thus improves the total system collection efficiency. Each individual imaging unit, including all electronic readouts, covered a 4.84 mm diameter footprint. Custom-sourced 2 mm diameter microscope lenses (1.5 - 3 mm working distance, 0.15 - 0.35 NA, 2-element sets) offer a central resolution of 8.8 - 27.9 μm depending upon the set working distance and FOV.

While the cumulative FOV of all four cameras provides a total imaging area of 45.9 mm^2^, we have currently configured the scope to image from a total area of ∼32 mm^2^ from across multiple regions of the dorsal cortex of a freely behaving mouse. The reduction in effective imaged FOV is due to the frontal micro-cameras imaging areas beyond the cranial window. Nevertheless, this is a nearly four-fold increase in imaging area as compared to existing miniaturized imaging devices (**Supplementary Fig. 1**), while minimizing the overall device weight and form factor to allow imaging in freely behaving animals (**Fig. 1H**) The micro-cameras located in the anterior locations cover most of the primary and secondary motor cortex regions along with parts of the somatosensory cortex in both hemispheres. The posterior micro-cameras were configured to image portions of the retrosplenial cortex, anterior parts of the primary visual cortex, and the association and the somatosensory cortices in both hemispheres.

### Optical Characterization

We performed a series of benchtop tests to evaluate and optimize the resolution and image FOV of each micro-camera in the mini-MCAM system. An image of the 1951 USAF test target (#58-198 Edmund optics) captured with a single micro-camera in the mini-MCAM is shown in **Figure 2A**. The imaging system can distinguish between both vertical and horizontal line pairs in group 6 element 5 which translates to a peak resolution measure of our system of 9.9 µm full-pitch resolution at a FOV of 4.5 X 2.55 mm. When configured to image a FOV of 3.56 X 1.99 mm we had the highest central full-pitch resolution of 8.8µm whereas in the corner of the FOV, resolution declined to 11 µm. This is on the lower end of the resolution range for similar miniaturized large FOV systems^17,28,29^ . Our focus was to image much larger FOVs at the tradeoff of slightly lower resolution. This alternate strategy has previously been used to successfully image large scale populations of neurons across large FOVs using low resolution optical imaging systems when the expression of the calcium indicators are sufficiently sparse^10–12,24,30^. To further quantify the resolution, we measured the light intensity variation across point sources by measuring 2 µm fluorescent beads (**Fig. 2D**). The resultant intensities across multiple beads were averaged to estimate a full width half maximum (FWHM) value of 7 ± 0.4 µm along the vertical axes and 14 ± 0.4 µm across the horizontal axes. At the corners, FWHM increases to 14 ± 0.4 µm along the horizontal and 21 ± 0.4 µm along the vertical directions. The increased FWHM at the corners is a result of the optical aberration that is inherent to the plastic micro lens objective that we are currently using. We then performed a wide-field imaging of a 400 µm thick coronal brain slice from a transgenic mouse specimen (Cux2-CRE-ERT2 x Ai162^30^) with sparse expression of GCaMP6s in layers 2/3 pyramidal neurons. Thus, the Mini-MCAM can discriminate between individual neurons expressing GCaMP6s (**Fig. 2F**).

**Figure 2:**
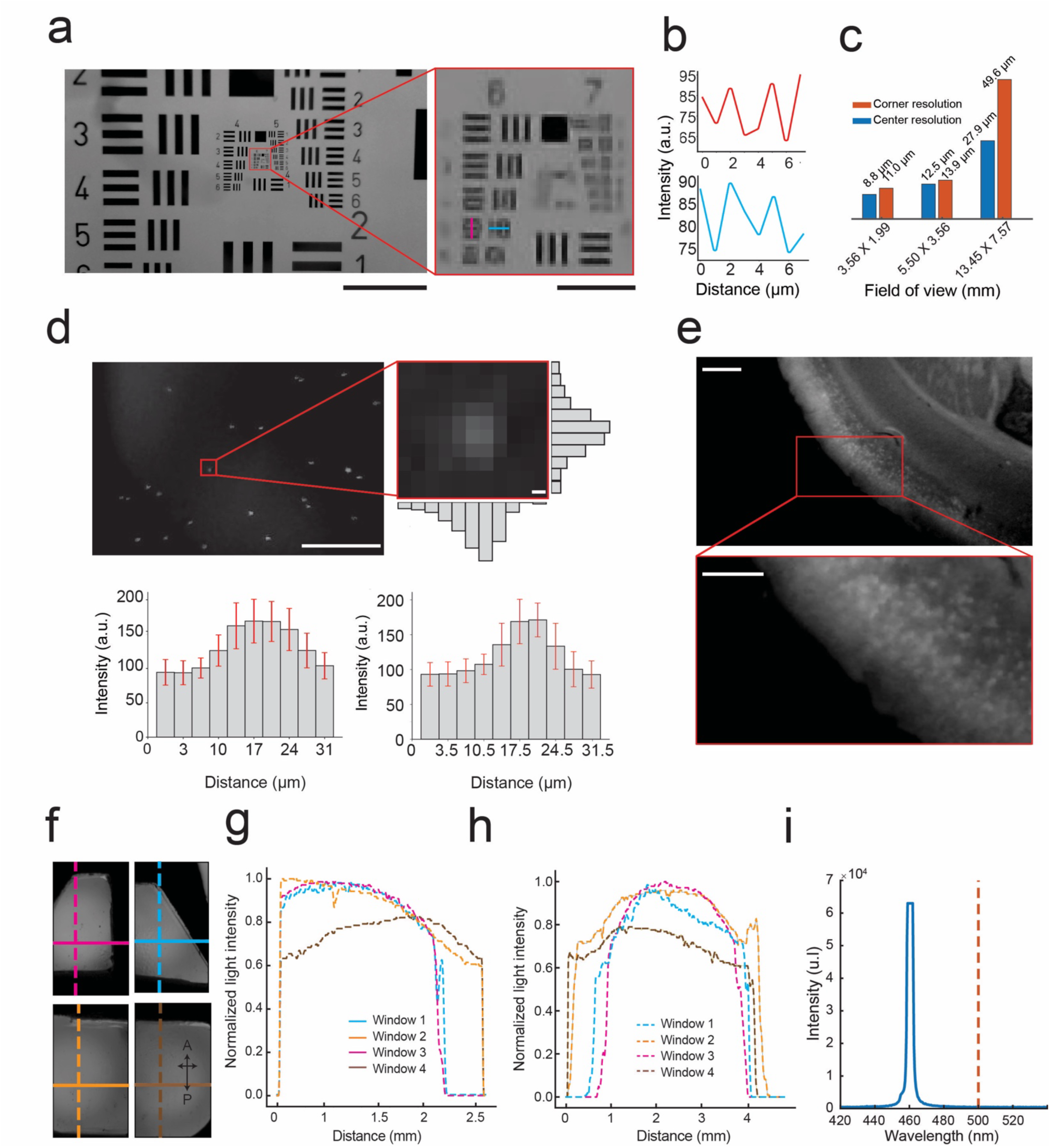
Optical performance of mini-MCAM: (a) *Left:* Image of the #58-198 Edmund optics USAF resolution test target captured by a camera in the mini-MCAM. Red square outlining group 6 and 7 elements. Scale bar indicates 1 mm. *Right:* Zoomed in view of group 6 and 7 elements from the image shown on the left. The magenta and cyan lines indicate the smallest elements that can be distinguished. Scale bar indicates 100 µm. (b) The intensity profiles measured across the magenta and cyan lines in the image shown in a. right. (c) A bar graph plot of center and corner resolutions as a function of image FOVs achieved at different focal lengths. (d) Image of 2µm fluorescent beads suspended in aqueous medium captured by a camera in the mini-MCAM. Scale bar indicates 2 mm. Right inset: light intensity variation across a single fluorescent bead. Bar graphs indicate the mean vertical and horizontal intensity variation (error bars show 1 standard deviation) across all microbeads in the field of view. Scale bar indicates 2 μm. (e) An image of a 400µm thick coronal brain slice from a Cux2-CRE-ERT2. x Ai162(TIT2L-GC6s-ICL-tTA2 mouse captured using the mini-MCAM camera; inset: zoomed in view of a small section showing individual cell bodies in layers 2/3 of the brain slice. Scale Bar indicates 2 mm. (f) Image of fluorescein dye infused agar phantom captured by the mini-MCAM through the multiplanar faceted cranial window. The colored lines indicate both vertical and horizontal sections along which the light intensity profiles were obtained. (g) Illumination profile measured across each horizontal section indicated in figure f. (h) Illumination profile measured across each vertical section indicated in figure f. (i) Light power spectrum for the blue excitation light source which shows peak power cut off below 500 nm.

We next evaluated the uniformity of illumination obtained across each FOV using the laser diode coupled optic fibers (**Fig. 2D**). Light is filtered at the source prior to coupling with the optic fiber, thus resulting in a clean narrow band illumination (461.3 ± 3.3 nm), (**Fig. 2G**). Fiber coupling also allowed much higher light power delivery as compared to LED illuminators. Each FOV has a dedicated fiber providing up to 6.7 mW/mm^2^ of light power. The tips of each optic fiber were cleaved and then polished using sandpaper (**See Methods**). This resulted in a uniform distribution of light throughout the FOV in both the anterior-posterior and medio-lateral directions, with light powers generally highest at the center of the FOV and reducing to a minimum of 20% at the periphery compared to the center (**Fig. 2H-J**).

### mini-MCAM enable cellular resolution imaging across the cortex

To demonstrate the *in vivo* imaging capabilities of the mini-MCAM, we imaged awake head-fixed mice running on a disk treadmill adapted from our previous work^5,8^ (**Fig. 3A**). We utilized a transgenic mouse line (Cux2-CRE-ERT2 x Ai162) that sparsely expresses GCaMP6s in layer 2/3 excitatory neurons throughout the cortex^11,13^. **Figure 3B** shows a composite image consisting of a single brightfield image of the entire imaging area under the multi-planar faceted cranial window with maximum intensity projection images of time series activity measurements from each of the four cameras overlaid. Zoomed in view of maximum intensity projection images from two of the posterior cameras are shown in **Figure 3C**.

**Figure 3:**
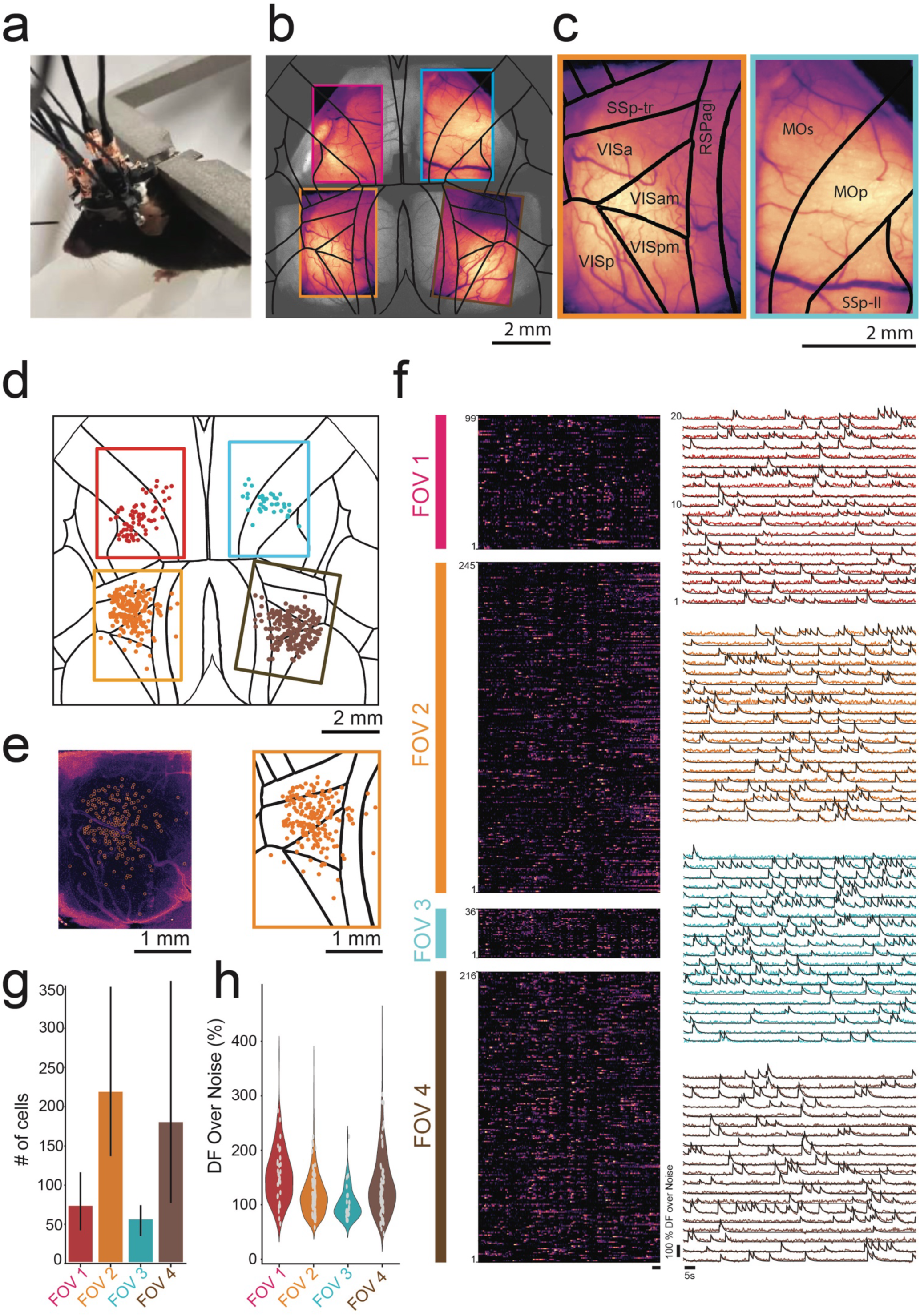
In vivo headfixed imaging using mini-MCAM: (a) Photograph of a head fixed mouse with attached mini-MCAM. (b) Maximum intensity projection (MIP) images from videos captured from the four cameras in the mini-MCAM overlaid over anatomically identical regions of the cortex in a widefield image captured from the same mouse. (c) Zoomed in view of the MIP images from the bottom left (orange outline), and top left (cyan outline) with the Allen CCF map overlaid. Abbreviations SSp-tr-primary somatosensory area-trunk, RSPagl - retrosplenial area agranular, VIS-Visual area (a- anterior, am- anteromedial, p- primary, pm- primary medial), MOs-secondary motor area, MOp- primary motor area, SSp-II - primary somatosensory *lower limb*. (d) Spatial coordinates of all cells with DFF > 20% recorded during spontaneous behavior in a head fixed mouse overlaid on the Allen CCF. Circles are color coded based on FOV they were recorded in. (e) *Left:* Background subtracted MIP image from the bottom right micro-camera shown in (b). white circles indicate detect cells shown in (d). *Right:* Zoomed in view of the same window from (d). (f) *Left:* Pseudo color normalized intensity plots of all cells recorded from each of the four FOVs indicated in (d). Scale bar indicates 5 seconds. *Right:* Raw DFF traces of 20 randomly selected cells in each FOV. (g) Bar plot of number of cells recorded in each FOV (n = 3 mice). (h) Violin plot of Average peak DFF measured in cells recorded in each FOV (n =3 mice).

Individual cells were readily discriminated even in the raw images captured by the micro-cameras (**Supplementary Video 1**). The raw images were corrected for large rigid motion artifacts and temporally downsampled to reduce processing time before applying spatial and temporal filtering. Since we are capturing slow Ca^2+^ transients (6 - 10Hz), we noticed a negligible loss of data fidelity by sub- sampling the imaging dataset at 15 Hz. These pre-processed videos were then analyzed using a cell activity extraction algorithm CNMFE^31,32^ (see **Methods**) extracts footprints of recorded cells (**Fig 3D** and **E**). In general, we found that the highest density of cells was found at the center of each FOV, possibly owing to degradation in the overall resolution in the peripheral regions as well as lower intensity of illumination from the optic fibers. We were able to extract Ca^2+^ activities from hundreds of cells in each FOV, spread across multiple brain regions from all three mice. Spontaneous activity in head-fixed mice were highly synchronized, with bands of high activity frequently appearing in the normalized Ca^2+^ activity traces correlated with motor activity on the treadmill (**Fig. 3F, left**). Examples of 20 randomly selected neurons in each window imaged in one mouse is shown in **Fig. 3F (right)**. Overall, across mice (n = 3), when limiting to cells with peak-to-noise ratio above 75 (**Fig. 3H**), we were able to capture 217 ± 107 (mean ± SD) neurons and 178 ± 143 neurons in each of the posterior windows, with 71 ± 40 neurons and 54 ± 19 neurons in each of the anterior windows where only part of the imaged FOV contained active brain regions. Compared to results in recent works^17,28,29^, the number of neurons recorded per unit area field of view is on the lower side which we believe is due to two reasons. Firstly, the endoscopic CMOS sensors used have reduced sensitivity. Secondly, the transgenic animals used for sparse expression have lower density of Ca^2+^ indicator positive cells. In the future, micro-cameras incorporating high sensitivity CMOS sensors could be integrated within the mini-MCAM. Further, alternate strategies for sparse and broad expression of Ca^2+^ indicators could also be explored^10,24^. Nonetheless, the distribution of DFF values that has been estimated for each putative cell region across all the mouse trials shows that most of the traces detected have an average peak DFF value between 50-200% (**Fig. 3H**).

### Imaging distributed cortical neural populations in freely behaving mice

Using the mini-MCAM, we next performed Ca^2+^ imaging in freely behaving mice. Mice head-mounted with mini-MCAM were allowed to spontaneously explore a custom-built linear track. To assist with signal transmission and delivery of illumination light to the mini-MCAM, we used a computer vision-guided motorized commutator^33^ that tracked the position and angular orientation of the mouse within the linear maze using real-time behavior imaging camera and adjusted the position and angular orientation of a slip-ring commutator. This real-time adjustment allowed us to alleviate any torsional stressors and allowed us to limit the length of the interface cables to < 1.5 ft (**Fig. 4A**). Mice readily explored the linear maze arena when head mounted with the mini-MCAM and completed multiple laps within recording sessions lasting 10 minutes (**Fig. 4D top**). Like headfixed recordings, we were able to observe single neuronal activities in the raw Ca^2+^ imaging videos (**Fig. 4 C, Supplementary Video 1**). Overall, we recorded from a total 492 neurons across all the fields of view. Onset of movement resulted in broad activation of neurons distributed throughout the cortex, consistent with observations in previous studies (**Fig. 4D)**^3,4,11^. Example Ca^2+^ activity traces from cells randomly selected from the normalized activity plots are shown in **Fig. 4E**. These experiments demonstrate the utility of the mini-MCAM for studying large scale neural activity across multiple brain regions while the mouse performs complex behaviors in a naturalistic setting.

**Figure 4:**
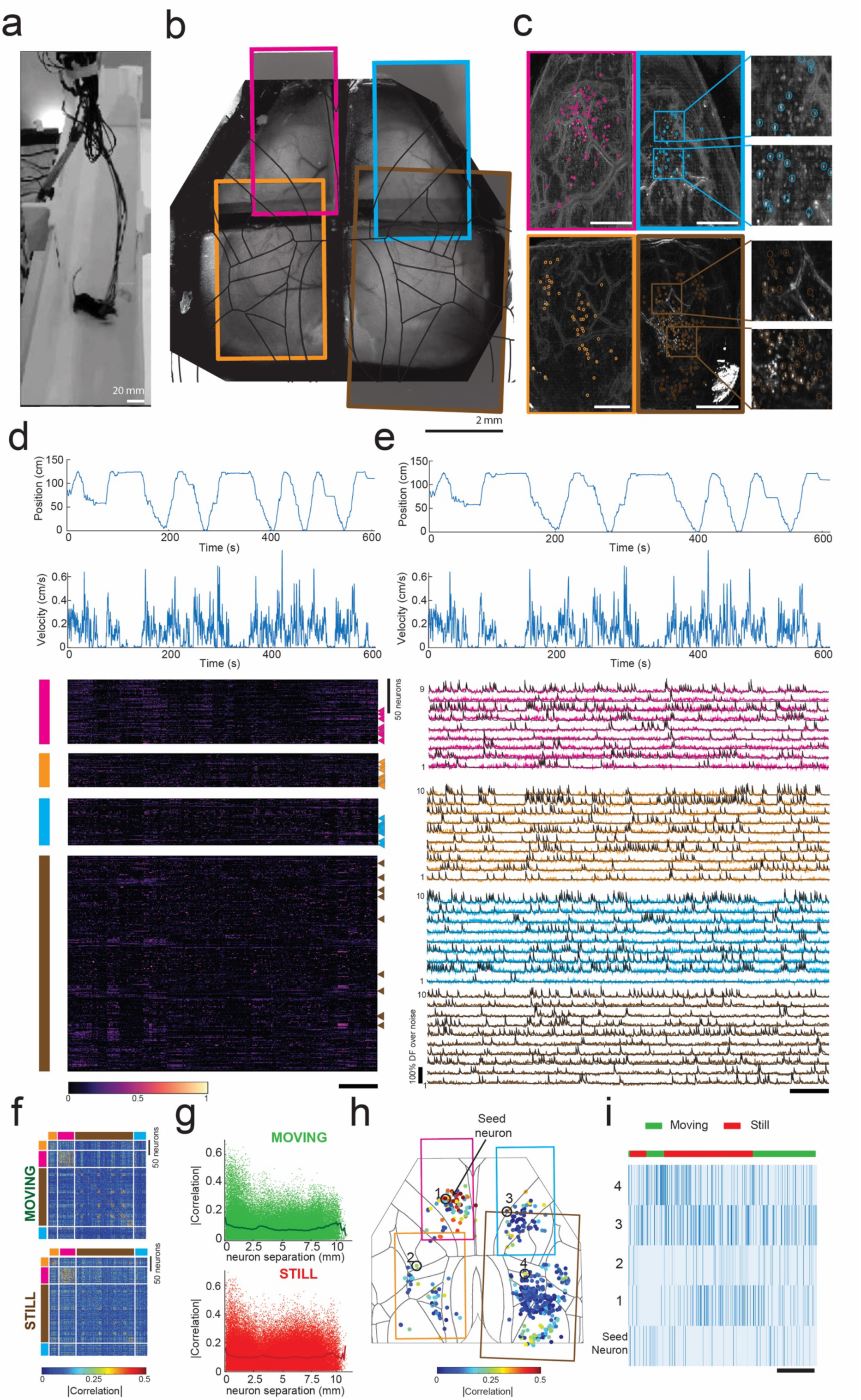
Cortex-wide mini-MCAM imaging in freely behaving animals: (a) Photograph of a freely moving mouse with mini-MCAM in a linear track arena. (b) A brightfield image of the mouse brain surface with the cranial window implant overlaid with the four FOVs captured using the mini-MCAM. Colored rectangles denote the position of the camera FOV. (c) Background subtracted max projection images for each FOV shown in (b). Circles indicate each cell detected using the cell extraction algorithm, insets. Scale bars indicate 1 mm. (d) The position and velocity x-direction aligned with the heatmap of the normalized z-scored DF/F traces for all cells detected in each window. colored vertical bars indicate FOV they were recorded from. triangles indicate the cells whose activity traces are shown in (e). Scale bar indicates 60 seconds. (e) Calcium activity traces from randomly selected neurons from each FOV. Scale bar indicates 60 s. (f) Spearman’s correlation coefficient matrices of cell-cell correlation during a 10-minute trial during epochs of movement and stillness. (g) Spearman’s correlations for each individual cell with respect to all cells recorded as a function of inter-neuron distance. (h) Spearman’s correlations for each individual cell with respect to a seed neuron denoted as an enlarged solid red circle. Colored rectangles indicate the borders of the imaged FOVs. (i) Raster plots of inferred spiking of the seed neuron and 4 randomly selected neurons indicate in (h). Scale bar indicates 25 s.

Simultaneous measurement of neuronal activity distributed throughout the cortex allowed us to evaluate the structure of neuron-neuron correlations during both periods of stillness and during movement. We computed the correlation magnitude (at zero lag) of the activity of each neuron with respect to all neurons recorded across the four FOVs (**Fig. 4F-I**). In general, neurons tended to be correlated to neurons located close to them, under both still and moving conditions (**Fig. 4G**). Evaluating the spatial distribution of cell-cell correlations, for one randomly selected neuron in the upper left FOV (**Fig. 4H**), we find clusters of cells proximal to the seed neuron with high correlation, but several examples of highly correlated neuron pairs distributed throughout the cortex were also observed (**Fig. 4I**).

## DISCUSSION

The goal of this study was to realize a miniaturized MCAM for imaging in freely behaving animals. The combined FOVs made available for optical imaging by the multiplanar faceted cranial windows offer several opportunities for very large-scale neural imaging and perturbation across the cortex with array optical neural devices, particularly for headfixed studies. First, arrays of high-performance large FOV imaging devices could be specifically designed, incorporating traditional CMOS sensors^26,34,35^, or those adapted for ultra-fast voltage imaging^36^ can be developed for multisite voltage imaging across multiple brain regions. Second, devices for patterned optogenetic stimulation of one FOV while imaging multiple other FOVs and combinations thereof can be designed in the future. While we attempted to maximize the area of imaging FOVs, while discriminating single-cell activities, the micro-cameras can be configured to both reduce FOV while increasing resolution, in the case of lower sensitivity imaging. Or increase FOV to simultaneously imaging multiple overlapping FOVs^37^, albeit at lower resolution. Thus, the mini-MCAM offers flexibility for multi-site studies that are difficult to achieve using single optical imaging systems.

Several other possibilities emerge with the use of arrayed micro-cameras. While we focused on covering most of the dorsal cortical surface, the array layout and positioning of each micro-camera can be reconfigured to allow simultaneous wide-FOV imaging across other regions - for instance, simultaneously imaging both the cortical and cerebellum via large cranial windows^38^. The relatively small footprint of the micro-cameras (<55 mm diameter) allows other modalities to be combined with cortical surface imaging. For instance, the micro-camera approach could be combined with GRIN lens-based miniscopes^35,39–41^ for deep brain cellular imaging simultaneously with multi-region cortical imaging. Further, the mini-MCAM can be particularly useful for very large FOV imaging across larger animal models such as rats and non-human primates such as marmosets cortex, where transgenic strategies for broad expression of calcium indicators are now emerging^42,43^ Polymer substrates used for defining the cranial windows can also be functionalized with flexible electronics for simultaneously recording surface field potentials^44,45^.

Compared to other miniaturized systems, the number of neurons we detected per unit area imaged is low. We hypothesize that this might be a result of multiple factors - the transgenic mice we used to express GCaMP result in Cre- recombinase production as a function of tamoxifen dosage. This may result in uneven expression of the Ca^2+^ indicator, as we previously observed^11^. This could be alleviated by exploring alternate strategies for sparse expression of Ca^2+^ indicators^10,12,24^. The off-the shelf microendoscope cameras adapted for fluorescence imaging have lower sensitivity as compared to traditional miniaturized imaging devices^18,26,34,36^. This results in lower SNR, thereby reducing the number of cells we can confidently annotate as cells. In the future, mini-MCAM imaging coupled with higher sensitivity calcium indicators that are significantly brighter than GCaMP6, such as GCaMP7^46^ and GCaMP8^47^ could result in a higher number of cells detected owing to higher SNR. In the current implementation, each micro-camera is attached. Finally, the mini-MCAM as used utilized the native electronic readouts of the endoscopes that were adapted for this application. This resulted in an overall increase in the weight of the device. In future iterations, custom serializer circuits that replace thick serial bus cable interfaces with thinner co-ax cables^48^ could be used to significantly reduce device weight and increase animal mobility.

## METHODS

### Multi-planar faceted cranial windows assembly

The cranial window frame was 3D printed with black resin (Black v4, Formlabs). The frame was partitioned into four windows using the horizontal and vertical cross bars. The experimenter fitted each of the four windows with transparent PET films (150 μm) that were laser cut to match the perimeter of each of the windows. First, the protective layers pre-bonded with the PET films were stripped using forceps. The exposed PET cut pieces were then bonded to the corresponding window frames using optical adhesives (NOA61, Thorlabs). The surface was then exposed to UV light (315-400 nm) for approximately 60 seconds to cure the optical adhesive and adhere the PET material to the frame. A thin layer of SU8 material (XP PrilElex SU-8 1.0, Micro Chem) was applied to the inner surface of the PET to stiffen the film and enhance its biocompatibility^49^. The SU8 coating was pre-baked at a ramping temperature from 60℃ to 90℃ for 1 min, then placed under 365 nm UV light to polymerize the monomer followed by post-baked at a ramping temperature from 60° C to 90° C. The head-plate was water-jet cut from a 0.016-inch-thick titanium sheet. The implant was then attached to the titanium head plate using the three fixing holes that were tapped using 0-80 hand taps. The head-plate design allowed for both head-fixation of the mouse and for attaching the Mini-MCAM using a form-locking mechanism. A custom 3D printed protective cap could be connected to the head-plate using two of the two mounting screw holes on the titanium head-plate to prevent dust and debris from accumulating on the cranial window surfaces when imaging was not being performed

### Mini-MCAM fabrication and assembly

#### Micro-camera assembly

Each micro-camera was built by adapting a micro-endoscope camera (TD-B209M3-76-01, MISUMI Inc.) for fluorescence imaging. The distance between the objective lens and the sensor was adjusted to achieve the desired resolution and FOV. A custom diced 4 mm diameter bandpass emission with pass band of 500-550 nm (ET525/50m, Chroma. Inc.,) was bonded on the objective lens using optical glue (NOA61, Thorlabs Inc.). Heat conductive Copper tape was wrapped around the micro-camera and emission filter to reinforce the structure. The camera with the emission filter was then placed in an aluminum heat sink tube with an inner diameter of 4 mm and an outer diameter of 4.5 mm, and silicon elastomer (Kwik-Sil^™^, World Precision Instruments) was used to seal the assembled micro-camera.

#### Illumination

A custom-built laser-coupled optical fiber bundle was used for illumination. A diode laser (465 nm, 2000 mW 2-Pin Blue Pigtailed Laser, CivilLaser) was installed in a lens holder (SM01, Thorlabs Inc.) The ends of each optic fiber in the bundle were polished first using SMA fiber polish kit (Thorlabs, CK01) to create a perpendicular polished surface facing the blue laser light. A bundle of 4 optic fibers was mounted within a custom fiber holder. a bandpass optical filter 440 nm – 480 nm (ET460/40 nm, Chroma. Inc.) was placed between the laser diode and tips of the optic fiber bundles for filtering of excitation light. The laser diode power was set to a constant voltage of 3.61 Volts, drawing an average current of 0.7 A. Under optimal conditions, the laser diode allowed us to achieve an illumination light intensity ranging from 3.3 mW to 4.2 mW at each optic fiber’s cleaved and polished ends.

#### Mini-MCAM assembly

The main housing of the mini-MCAM was 3D printed out of black resin (Black v4, Formlabs). The housing consisted of four slots for the insertion of the imaging cameras, each with an inner diameter of 4.6 mm. Assembled micro-cameras were inserted into one of four camera slots in the main housing. Corresponding illuminator fibers were inserted into the illumination sleeves and bonded using epoxy.

### Benchtop testing of the mini-MCAM

#### Resolution testing

A 1951 USAF test target (#58-198 Edmund optics) was used to measure one set of micro-cameras’ peak central and corner resolutions in the mini-MCAM array. The resolution test target was placed on a custom 3D-printed platform below the micro-camera and a white light source was used to back illuminate the test target. A custom 3D printed fixture mounted on a translation stage (PT3XYZ stage, Thorlabs) was used to house the micro-camera, which allowed adjustments to the camera’s working distance and lateral position with respect to the test target.

#### Characterizing Illumination profile

Custom 3D printed cavity was filled with fluorescein dye-infused with 3% agar gel (10% v/v, catalog no. F2456; Sigma Aldrich). The cranial implant was then placed within a mold made on the open surface of the cavity that leveled the top surface of the implant with the top surface of the cavity and filled up the entire bottom transparent PET window surface with fluorescent gel. The titanium head plate was screwed on top of the cranial implant, and the Mini-MCAM was then attached to the head plate to emulate similar conditions as that of an in-vivo experiment. We then simultaneously recorded from all four windows at 30 Hz. The illumination from the optic fiber was modulated to avoid pixel saturation in the recorded FOVs. The video dataset was then analyzed using a custom Python script.

#### µ-bead imaging test

10 µl stock solution of 2 µm fluorescent beads (Lot No. 189907, Thermo Scientific) was diluted by mixing into 10 ml of distilled water. The diluted µ-bead solution was then stirred with an orbital shaker for 1.5 hours to create a uniform suspension. 1 µl of the suspended solution was mixed with 1mL of ethanol for the final dilution and placed back in the orbital shaker for 30 minutes. A single drop of the final dilution was spread uniformly on a glass slide with a coverslip placed on top to spread the micro-bead sample. The slide was then imaged using one of the micro-camera in the mini-MCAM array. We exposed the µbead glass slide-to-bottom illumination using a custom LED illumination light source with a peak excitation light wavelength cut off at 470 nm.

#### Imaging fixed brain slices

400 µm thick coronal brain slices from double transgenic mice (Cux2-CRE-ERT2xAi162) expressing GCaMP6s sparse populations of layer 2-3 pyramidal neurons were imaged using the micro cameras. The optic fiber illumination used in the mini-MCAM was used for illuminating the brain slice for consistency.

#### Surgical preparation

All animal experiments were approved by the University of Minnesota Institutional Animal Care and Use Committee (IACUC). Double transgenic mice that resulted from cross-breeding Cux2-Cre-ERT2 with Ai163-GCaMP6^50^ were used in the mice. Mice were housed in a 14hr light/10hr dark cycle in rooms maintained at 20-23 °C and 30-70% relative humidity. Mice had ad libitum access to food and water. Mice were injected with 75 mg/kg of Tamoxifen 5 days prior to surgery to stimulate expression of GCaMP sparsely in layer 2/3 pyramidal neurons in the cortex^30^. The surgical methodology for implanting the multi-planar faceted cranial windows was adapted from our previous studies^25,30,51^. Briefly, mice were administered 2 mg/kg of sustained-release buprenorphine (Buprenorphine SR-LAB, ZooPharm) and 2 mg/kg of meloxicam for analgesia and preventing brain inflammation respectively, 40-60 minutes prior to surgery.

Mice were placed in a chamber and anesthetized with 1-5% isoflurane anesthesia in oxygen. Eye ointment (Puralube, Dechra Veterinary Products) was applied to the eyes. The scalp was shaved and cleaned. Mice were then transferred and fixed in a stereotaxic (900LS, Kopf), and the scalp was sterilized by repeatedly scrubbing Betadine and 70% Ethanol solution (3 times). Next, the scalp was removed using surgical scissors. The tissue and fat under the scalp were subsequently cleared using a micro curette (# 10080-05; Fine Science Tools). The partial temporalis muscle wrapping around the skull was carefully removed using a scalpel. The frame of the cranial window implant was placed on the skull, and the outline of the craniotomy was marked using a surgical marker and subsequently scored using a curette. A high-speed dental drill was used to perform the craniotomy. All cranial windows were implanted symmetrical about the midline suture, with the anterior boundary of the implant placed 4 mm anterior to Bregma and the posterior boundary placed 3.9 mm posterior to Bregma. The brain was soaked in gel foam soaked in sterile saline until bleeding from the craniotomy stopped. The skull surrounding the craniotomy was cleaned using a cotton tip applicator. The cranial window was placed gently on the exposed brain, and surgical adhesive (Vetbond, 3M) was applied around the edges of the cranial window frame to adhere the implant to the skull. After the adhesive was cured, the Titanium head plate was fastened on the implant with #0-80 screws. The implant was cemented to the skull using opaque dental cement (Metabond, Parkell Inc.). Mice were transferred onto a heating pad to recover from the surgery and then transferred to a clean cage after they were full ambulatory. Mice were allowed to recover for 6-7 days post-surgery prior to any behavioral or imaging studies.

### Headfixed imaging using mini-mCAM

#### Head-fixed acclimatization

Mice were allowed to recover up to 1 week after surgery before acclimatizing to the head-fixation apparatus. Animals were head-fixed to a custom-built headstage^5^ for 5-15 minutes every day for 5-7 days until they were comfortable spontaneously running the treadmill. The experiment design is adapted from our previous study of ECoG recording^45^

#### Mini-MCAM focusing

Once the mice were head-fixed to the custom head-stage, each cranial window was cleared of debris using a cotton Q tip dipped in deionized water. Each micro-camera was manually focused by adjusting the position of the micro-cameras within the cylindrical channel in the integrated camera housing until distinct single-cell outlines were observed in the live streams of each FOV using OBS studio software. Once optimal focus was achieved, the focus lock set screw was tightened to maintain the relative position between the micro-camera and the bottom camera housing. Hot melting glue was additionally applied around the micro camera to reinforce the camera further and minimize motion artifacts during imaging.

### Freely behaving imaging using mini-MCAM

The animal was acclimatized to the behavior arena over seven days prior to imaging. The animals were first handled in their home cages for 5-15 minutes for the first three days. Mice were next allowed to explore the open maze for 5-15 minutes while being head-mounted with a dummy mini-MCAM scope to acclimatize them to the weight of the scope. Prior to freely behaving imaging experiments, the mouse was attached to a custom head-restraint setup^5^, and the mini-MCAM was attached to the headpost. Individual micro-cameras were adjusted to focus on a layer of cells ∼150 µm below the surface of the brain focus. After reaching the best focus for each FOV, hot glue was used around each micro-camera to fix the position of each camera. On the day of the imaging experiment, the mice were introduced to the middle of the linear maze measuring 4’ x 6”. The walls of the linear maze were laser cut from 1/8” white opaque acrylic sheets. Two wide-angle cameras (Logitech C90 pro-HD Webcams) were fixed at elevated points to image the behavior of the mice within the arena. Each camera covered one-half of the line maze environment, and data acquisition was synchronized with the mini-MCAM using the OBS studio software platform. Datalinks from the mini-MCAM were routed through a custom-built active motorized commutator that utilized real-time markerless tracking of the mouse to detect heading orientation and periodically corrected for accumulated twisting of the cables^33^. Each imaging session lasted 10 minutes.

### Imaging Data analysis

Ca^2+^ trace datasets were simultaneously recorded at 30Hz using the OBS studio software (Windows v. 30.1.2) on a large canvas of size 2560 x 1500 pixels for the headfixed trials and a canvas of size 4000 x 1440 pixels for the freely behaving trials. Each of the four mini-MCAM camera sensors occupied a pixel footprint of 1280 x 720 on the large canvas for both behavior assay recordings. In addition to the mini-MCAM camera feeds, the freely behaving datasets also captured frames from two behavior cameras, occupying a pixel space of 1440 x 720 pixels on the final canvas. The experimenter then saved the recordings in an mp4 format with a H.264 video encoder. The output from OBS studio software was dynamically cropped using MATLAB to generate separate video files for each FOV. Python script was then utilized to load these video files and save them as individual TIFF stacks. Custom python code was then utilized to perform rigid motion correction, and to apply spatial and temporal filtering as recommended in the Inscopix-CNMFE^53^ workflow guide. CNMFE^54^ was then utilized to perform cell extraction.

The image stack was down-sampled to 15Hz by accessing every other frame in the TIFF file. Subpixel rigid motion correction was applied to the image stack using the phase cross-correlation algorithm ^52^ provided in the scikit-image Python package for image registration. We set the maximum allowed shift to be 30 by 30 pixels in each dimension to avoid spurious correlations. A temporal median filter and a savitzky-golay filter (Scipy multidimensional image processing package^53^) was applied to eliminate low-frequency background noise (< 3Hz). Finally, a baseline filter was used to remove low-frequency fluctuations in the temporal signal. A spatial gaussian low-pass filter (σ=0.5) and a uniform high-pass filter with a kernel size of 13 were applied to smooth out the high-frequency spatial noise and to remove background fluorescence respectively.

Cell extraction was performed with Inscopix’s open-sourced CNMFE implementation^54,55^. We set the mean cell diameter and the ring size factor to be 5 pixels and 2 respectively and found that default options for the remaining parameters gave the best results for cell extraction. The raw traces and footprints of the putative cells extracted from the algorithm were then further filtered by thresholding based on peak-to-noise ratio (PNR) between 75 and 175, and area thresholding to remove regions of interest with abnormally large cell footprints.

## ACKNOWLEDGEMENTS

We acknowledge the staff at Research Animal Resources, University of Minnesota, for animal care and housing; Leila Ghanbari for the initial exploration of the mini-MCAM concept. Funding sources: UMN Mechanical Engineering Department, MnDRIVE RSAM, National Institutes of Health (NIH) grant R01NS11128, and BRAIN Initiative grants RF1NS113287, SBK and RH; P30DA048742 and RF1NS126044 SBK.

## AUTHOR CONTRIBUTIONS

JH, AC contributed equally to the manuscript. JH, RFH, RH and SBK conceptualized the mini-MCAM. JH, RFH, and DAS designed and built the mini-MCAM hardware. JH, AC, IO and DAS performed benchtop testing and in vivo experiments. AC, DAS, RP and MF analyzed data and visualized data. KS, EK assisted with surgeries, implant preparation and behavior experiments. AC and SBK wrote the original manuscript draft. All authors reviewed and edited the manuscript. RH and SBK supervised all aspects of this work. RH and SBK acquired funding.

---

**SUPPLEMENTARY VIDEO 1: Pan Cortical Ca^2+^ imaging during spontaneous headfixed behavior: *Left:*** Raw Ca^2+^ videos of each FOV overlaid on the Allen reference atlas. ***Center:*** Highlighting single neuron activities in insets highlighted in whole FOVs in the left. ***Right:*** Normalized Ca^2+^ activity of each neuron detected, with circles representing the location of the cells in the cortex. Video rendered at 2X real time speed.

**SUPPLEMENTARY VIDEO 1: Pan Cortical cellular imaging in a freely behaving mouse: *Left:*** Position tracking of a freely behaving mouse with a headmouted mini-MCAM in a linear track arena when imaged. ***Center, top:*** Background subtracted videos of two FOVs imaged along with insets highlighted a smaller area within the FOVs. ***Center, bottom:*** Normalized Ca^2+^ activity of each neuron detected, with circles representing the location of the cells in the cortex. ***Right:*** Normalized Ca^2+^ activity traces of all neurons in the FOVs imaged. Video rendered at 1.5X real time speed.

## SUPPLEMENTARY FIGURE

**Supplementary Figure 1:**
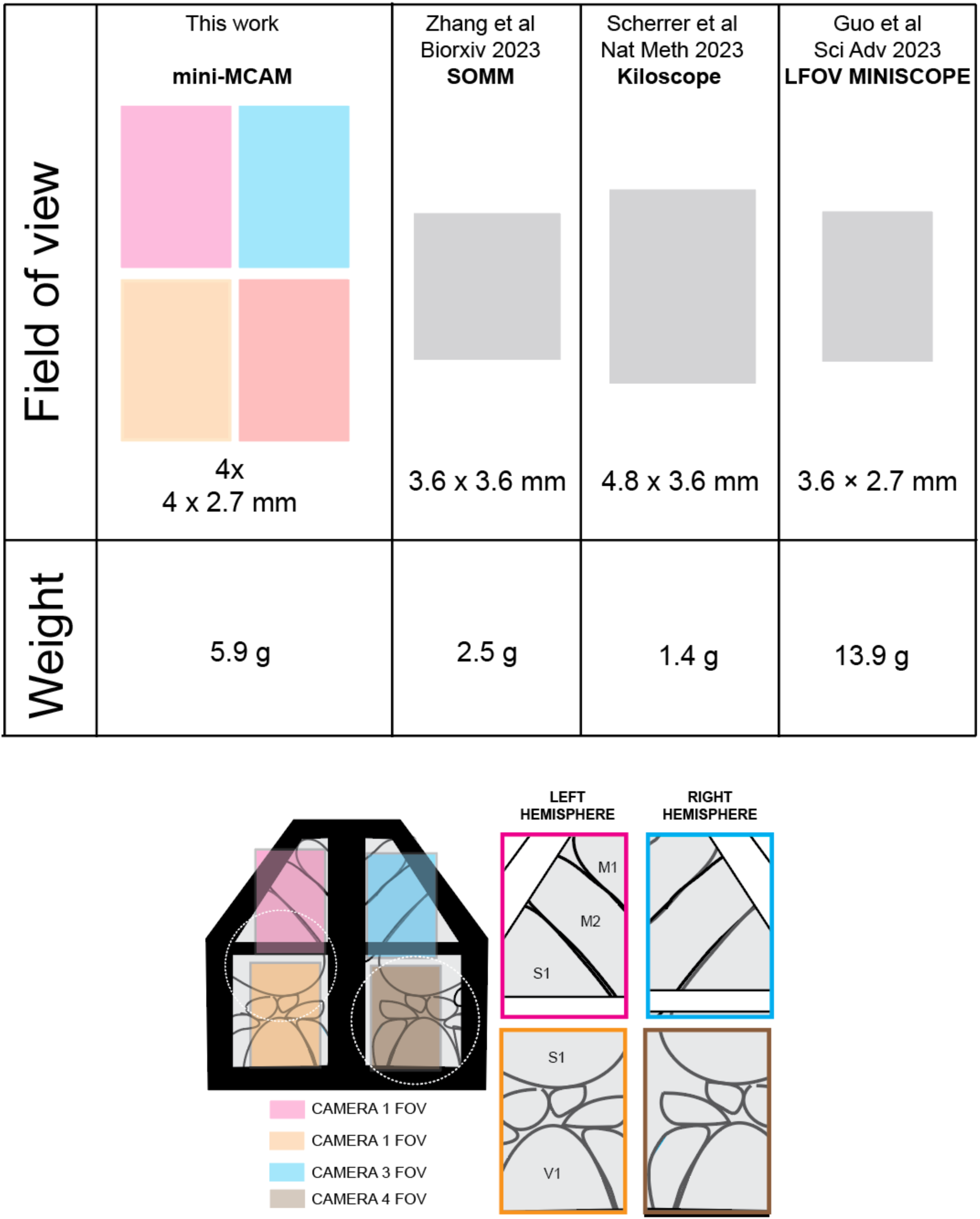
Comparison of the mini-MCAM cumulative FOV to existing large FOV miniaturized microscopes^1,2,3^.

